# Description of the sound diversity of two species of tuco-tucos (*Ctenomys torquatus* and *Ctenomys lami*) in natural environment

**DOI:** 10.1101/2021.06.01.446534

**Authors:** Keila C. Zaché, Lucas Machado Silveira, Gabriel Francescoli, Thales Renato Ochotorena de Freitas

## Abstract

Sound signals can travel through long distances, becoming an important communication channel between animals that need to establish contact beyond the visual form. They can then be considered a relevant form of communication between species living in the underground environment. *Ctenomys torquatus* and *Ctenomys lami* are solitary subterranean rodents, thus demanding an improvement of the communicative channels, especially in territorial defense actions and meeting potential reproductive partners. This work was aimed to describe the variability of acoustic signals emitted by *C. torquatus* and *C. lami* by analyzing the physical-morphological characteristics of the signals. The study was carried out in two populations, one of each species and was selected 14 individuals of *C. torquatus* and 15 *C. lami.* The acoustic signals were recorded in a natural environment, obtaining the sounds straight from the animal tunnels. A total of 1,380 signals were captured and analyzed, 786 from *C. torquatus* and 594 from *C. lami.* It was possible to characterize 5 different types of signals, emitted by both species. Most of the analyzed sequences presented low frequency, and many of these calls exhibited characteristics of long-range signals. It was verified a sharing of sound signals in both species, as regarding the acoustic parameters as the morphology of the analyzed spectrograms. For the first time, it was possible to have access to sound data emitted by direct subterranean rodents from their tunnels in the natural environment.

## Introduction

In order to exchange information, animals use different types of communication channels, such as chemical, visual, tactile and auditory signals [1]. According to Freeberg et al. [2] a greater communicative complexity is required of individuals living in more socially complex groups because the diversity of social systems correlates with the number of individuals who relate to each other, the frequency within which interactions occur, the social role of the individuals and the variability in the types of interactions. In heterogeneous communication systems, it is expected to happen a structuring of the acoustic elements, with a functional acting and a high amount of information [2].

Because of its plasticity, acoustic communication is considered very efficient [3], allowing in most vertebrates the composition of a sound signal that presents a wide variety of time, intensity and frequency variables, and this diversity may be related to multiple conditions [1]. A signal emitted by an animal may contain information about its social status, motivational status, and also the identity of the emitters, it may change according to behavioral contexts and environmental situations [1,4–7].

Subterranean rodents are naturally distributed on all continents, with the except for Australia and Antarctica, and they represent a unique group among mammals because they show different attributes due to their habitat [8,9]. The biotope where subterranean rodents are inserted is composed of a structure that is generally sealed, with a small diameter, much improved, and has a constant microclimate [9,10]. Over time, these characteristics may have acted in the development of the sensory systems of these animals [8,10,11]. In South America, there are two families of fossorial caviomorph rodents represented by Octodontidae and Ctenomyidae [9,12].

From the Ctenomyidae family, the species of the genus *Ctenomys* went through a process of fast speciation during the Pleistocene [13] showing high variability at the genetic level, expressing a wide degree in the speciation process [12,13]. They occupy a diversity of habitats, from drier fields to forest areas [9]. Most species have a solitary way of life, are typically territorial, structured in small semi-isolated populations with fragmented distribution, and low adult vagility [9,12–16].

There is wide use of the plurality of communication channels in these animals. Chemical and tactile signals are closely linked to the construction, use, and marking of burrows [17,18]. Acoustic signals have a fundamental role in inter-individual communication by emitting short or long-range signals, divided into vocal and non-vocal emissions [19,20]. Among the non-vocal signals are tooth beats and vibrational emissions [17,21]. Studies describe the presence of seismic communications in some subterranean rodents [22], but these emissions have not yet been reported in *Ctenomys*.

Vocal signals may be able to travel long distances and be compatible with underground life because they are required communicative peculiarities in this environment [23,24] since most subterranean animals perform a great part of their vital activities within tunnels [9,16]. One of the functions of long-range signals is the expression of territoriality, communicating the presence of the individual who owns a particular territory [19,20,25]. This type of signal could also act as a regulator of clashes between individuals and be significant in finding potential mates for reproduction [26].

About 69 species are described for the genus *Ctenomys* [27,28], and little is known about the acoustic repertoire of these species. Most sound data were obtained from laboratory work [19,20,25,29,30], and the scarce free-living data are from signals that can be heard and recorded outside their burrows [17,26,31]. There are no studies in the literature on captured vocalizations within the burrows of these animals in a natural environment.

*Ctenomys*’ vocalizations were studied in detail in only three species, the talas tuco: *Ctenomys talarum* [20,29], the Pearson’s tuco-tuco: *Ctenomys pearsoni* [19,26,30,31] and Anillaco tuco-tuco: *Ctenomys sp* [25,32]. In addition, Francescoli & Quirici [33] showed two possible vocal patterns that could be shared with other species of *Ctenomys.*

The species *Ctenomys torquatus* Lichtenstein 1830 is widely distributed, inhabiting the Pampa region, from the central region of Rio Grande do Sul to northern Uruguay [34] and the species *Ctenomys lami* Freitas 2001 is endemic to the state of Rio Grande do Sul, Brazil, occurring in a narrow range of fields in the region known as Coxilha das Lombas [35].

*C. torquatus* and *C. lami,* the two species focused on this study, live in regions that have suffered the effects of anthropization, transforming their areas of life into large extensions of reforestation, highways, and for *C. torquatus*, its geographic distribution in Brazil coincides with the activities of coal mining [36]. It is known that propagation of sound may suffer interference from anthropogenic noises, environmental temperature, humidity, and types of vegetation, therefore, the transfer of acoustic signals between individuals of these species may require modifications or even be compromised [1].

This work aimed to describe the variability of the main vocal signals emitted in wildlife by *C. torquatus* and *C. lami,* belonging to the same monophyletic group, *torquatus* [37], investigating the physical-morphological characteristics of these emissions. The vocal signals of *C. lami* have never been reported before, and *C. torquatus* was only superficially cited in a comparative work with other *Ctenomys* species [33]. Investigating the variability of the signals used by these animals would allow future comparison with the acoustic signals described for other *Ctenomys* species. Besides detailing the main sound attributes used by these two species, which would help studies on possible sound impacts caused by exposure and overlap of noises from anthropogenic activities, since sound signals are fundamental for the development and maintenance of these species.

## Methods

### Study field

The study was conducted in two populations, one of each species, located in different areas in the state of Rio Grande do Sul, Brazil. For the species *C. torquatus* the selected population was located in an area bordering the Ecological Station of Taim (32° 32’19.4 “S 52° 32’19.5” W), near the city of Rio Grande – RS – Brazil. For *C. lami,* the studied population was the one located inside the Biological Reserve José Lutzenberger Lami (30° 14’09.5”S 51° 05’45.3”W), situated in Porto Alegre – RS – Brazil.

The sampling period was ten days for each species. *C. torquatus* was sampled in June of 2017 (late autumn) and *C. lami* in October and December of 2017 (mid-spring).

### Burrows Selection

The burrows selected were the ones that presented recent activities. Recognition of recent activity occurs through the visualization of fresh sand mounds that are pushed out of the burrows during cleaning and foraging activities [9,37,38]. A minimum spacing of 15 meters was maintained between the selected burrows, considering the direction of the burrow tunnel gallery, to avoid capturing the same animal. After choosing the burrows, the trapping process was carried out.

### Trapping and capture process

The purpose of the use of traps was to capture the animals for weighing and sex identification. The capture technique consisted of using Oneida-Victor tramp traps, number zero, denveloped by rubberized material to avoid injuring the captured animal. These traps were introduced into the burrows and signaled with numbered flags, being reviewed every ten minutes. The traps remained in the selected burrows until the animals were captured or for a maximum of 40 minutes.

The project was licensed by SISBIO / ICMBIO, number 58477, and was approved by the Animal Ethics Committee of the Federal University of Rio Grande do Sul, number 33289, to conduct the research.

### Identification and selection of animals

After capture, the individuals were weighed and visually sexed [37,39–42], after performing these procedures, the animals were returned to the burrow from where they were captured. When the animal was not captured, but the burrow got closed, indicating that there was at least one individual inside the tunnel gallery, this individual was considered as unidentified. All animals had the position of their burrows registered using a GPS (Garmin Vista^®^).

Twenty-four *C. torquatus* individuals were selected: 3 males, 8 females, and 13 unidentified. Of these 24, it was possible to obtain vocalization from 14: 6 females, 2 males, and 6 unidentified. In *C. lami* 15 animals were selected, 3 females, 1 pup, which was not able to identify the sex, and 11 unidentified individuals. It was possible to obtain sound signals from all individuals.

### Data collection

The recordings were made with the aid of a Rode NTG-2 microphone connected to a Tascam DR-40 Digital Sound Recorder. To protect the microphone from possible damage caused by the animals, it was developed with the Mechanics Sector of the Institute of Physics of the Federal University of Rio Grande do Sul. This cylindrical apparatus that worked as a microphone protective capsule. A test was performed to verify if the cylindrical apparatus could modify the reception of the signals. For this test recordings were made within a burrow using both the microphone by itself and the microphone coupled with the cylindrical apparatus for subsequent comparison of the captured sounds. No significant change in signal reception was found, according to the sound frequencies captured.

The insertion of the microphone into the burrows occurred first with the removal of the sand deposited by the animal at its entrance, then was located in the main tunnel where the microphone was positioned. A minimum period of 24 hours was considered when the animals were captured and returned to their burrows so that the individuals could acclimate before the recording began.

A sampling effort of 65 hours of sound data collection for both species was made. For *C. torquatus,* 39 hours were recorded, and for *C. lami* were 26 hours. The individual recording time was at least 40 minutes.

### Sonographic analysis

The sonographic analysis was performed using Raven Pro software version 1.5 [43]. The spectrographic configurations were set up prioritizing the resolution and the easy detection of vocalizations, as follows: Window: Hann, 312 samples, 3 dB bandwidth-filter: 203 Hz; Time grid: overlap: 50%, hop size: 156 samples; Frequency grid: DFT size: 512 samples, grid spacing: 86.1 Hz.

The terminologies adopted in this paper to describe and identify the sound signals were note and phrase [44]. The note is a single sound unit with no interval, although the set of notes with silent intervals between them was called phrase (Fig 1). The acoustic parameters analyzed were: phrase size or note size, number of notes in the signal, dominant frequency, maximum frequency, and energy in the established sound sequences.

**Fig 1.**
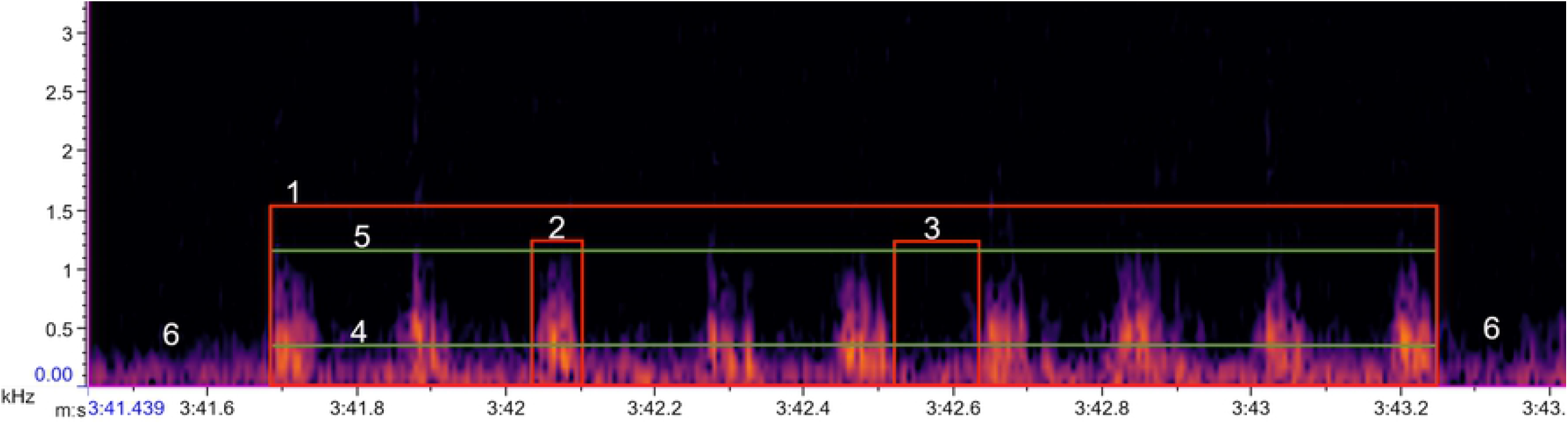
Spectrogram and acoustic parameters of a *C. lami vocalization.* The acoustic parameters are represented in red rectangles for (1) a sequential type phrase, (2) a note of sequential phrase and (3) the space between the notes. The green lines are (4) dominant frequency and (5) maximum frequency, on the outside of the marking is (6) background noise.

Phrases were manually selected according to the time and frequency limits arranged on the acoustic signals. The phrase length is the duration in seconds of the signal selection. The number of notes was counted visually and manually. The maximum frequency value coincides with the limit value of the phrase on the frequency axis. However, dominant frequency values and energy values were automatically generated according to specific formulas based on the average magnitude within the selected spectrum [45] (Fig 1).

## Results

From an auditory and visual investigation, it was possible to identify five different types of signals in *C. torquatus* and *C. lami*: mono, drummed, tuc, sequential and squeal (Fig 2), and a new nomenclature was made for the signals that not yet had been identified due to the absence of sound data for the species studied. A total of 1,380 signals were recorded and analyzed, 786 for *C. torquatus,* and 594 for *C. lami.* In *C. torquatus* the vocalizations were emitted by females, males, and unidentified individuals (Table 1), while in *C. lami,* the signals were emitted by females and unidentified individuals (Table 2). For all detected signals were measured the mean, standard deviation, and range (minimum and maximum values) of all acoustic parameters analyzed (Table 3).

**Fig 2.**
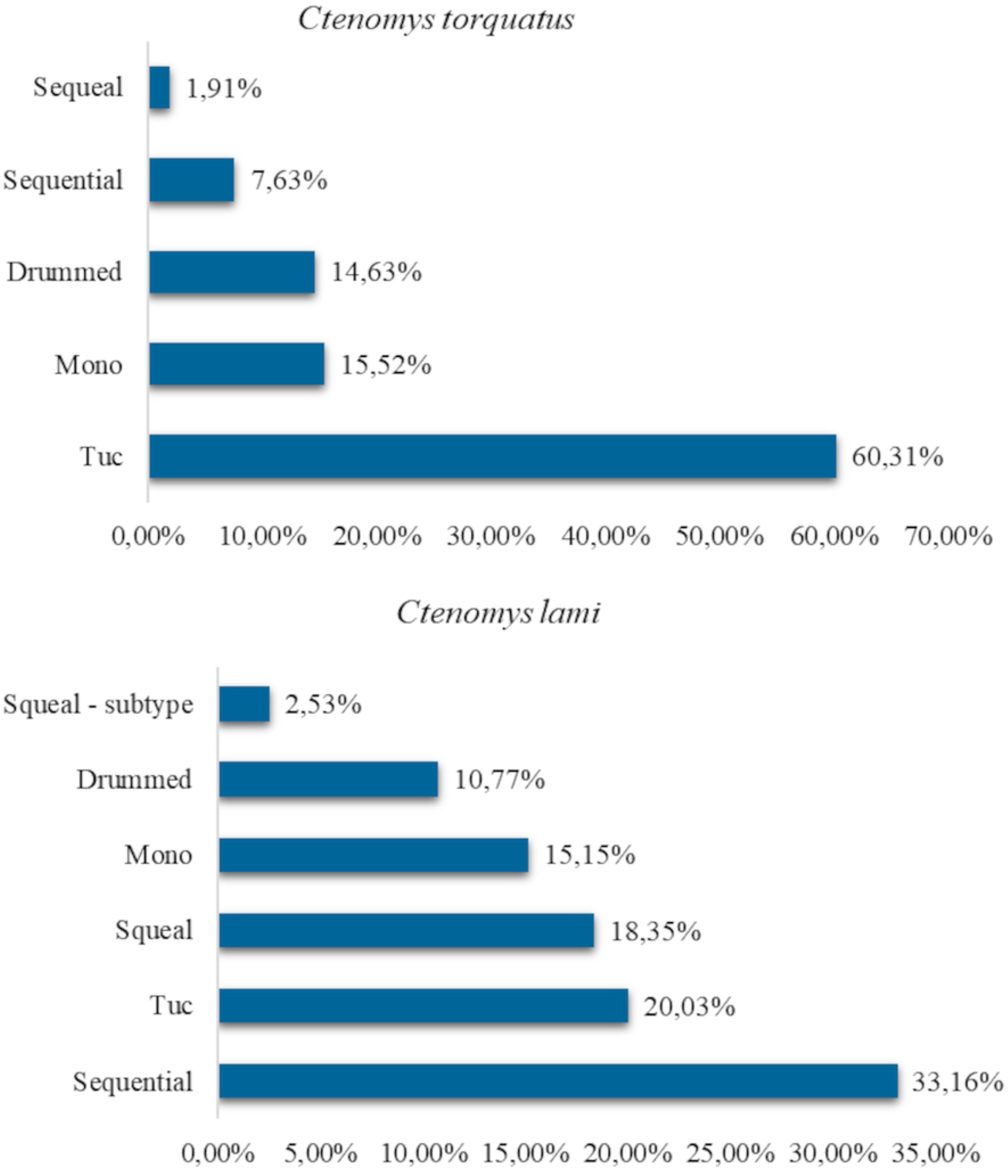
Signal types found in *C. torquatus* and *C. lami* populations. Percentage values of signals emitted by species, related from the least frequent to the most frequent signal.

**Table 1.** Total signals emitted by *C. torquatus* according to the sex of the individuals.

| Signal | Female | Male | Unidentified |
| --- | --- | --- | --- |
| **Mono** | 42 | 14 | 66 |
| **Drummed** | 51 | 4 | 60 |
| **Tuc** | 169 | 57 | 248 |
| **Sequential** | 51 | 4 | 5 |
| **Squeal** | 6 | 1 | 8 |

**Table 2.**
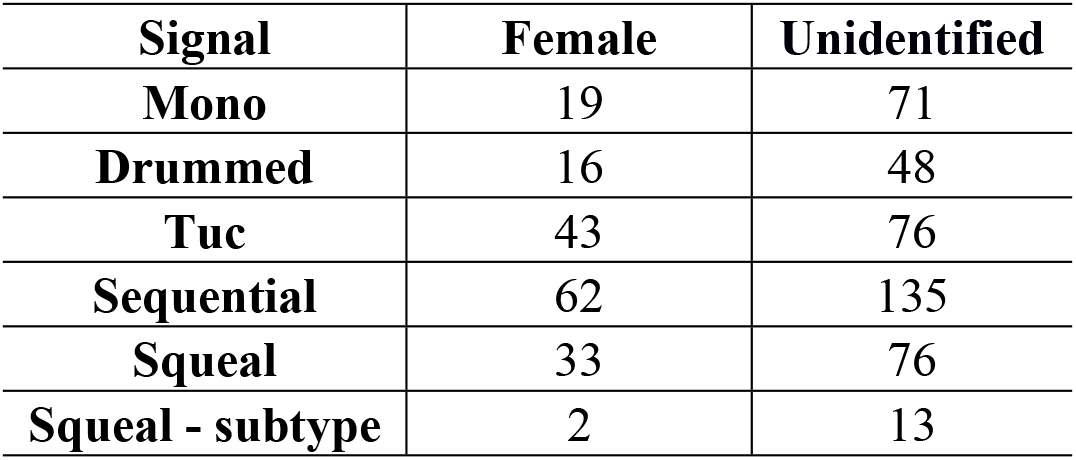
Total signals emitted by *C. lami* according to the sex of the individuals.

**Table 3.** Measurements (mean, standard deviation, minimum values and maximum values) of the acoustic parameters of the five described signals.

| Signal type | Species | Time (seconds) | Units | Frequency - kHz |  | Decibels - dB |
| --- | --- | --- | --- | --- | --- | --- |
|  |  | Signal length | Number of notes | Dominant | Maximum | Energy |
| Mono | *Ctenomys torquatus* | 0.030 (0.024) | 1 (0) | 0.460 (0.125) | 1.544 (0.672) | 71.766 (6.577) |
|  |  | 0.009 - 0.248 | 1-1 | 0.172 - 1.034 | 0.729 - 5.064 | 55.100 - 87.700 |
|  | *Ctenomys lami* | 0.067 (0.033) | 1 (0) | 0.215 (0.143) | 1.167 (0.590) | 69.152 (10.948) |
|  |  | 0.021 - 0.198 | 1-1 | 0.086 - 0.689 | 0.513 - 4.111 | 53.200 - 103.300 |
| Tuc | *Ctenomys torquatus* | 0.273 (0.129) | 3.152 (1.612) | 0.220 (0.091) | 0.771 (0.591) | 73.982 (12.141) |
|  |  | 0.071 - 0.886 | 2- 12 | 0.086 - 0.603 | 0.311 - 5.923 | 56.000 - 119.200 |
|  | *Ctenomys lami* | 0.347 (0.151) | 3.697 (1.482) | 0.219 (0.113) | 1.860 (1.503) | 88.816 (12.671) |
|  |  | 0.088 - 0.892 | 2- 12 | 0.086 - 0.603 | 0.340 - 7.372 | 57.100 - 119.100 |
| Drummed | *Ctenomys torquatus* | 0.326 (0.203) | 2.791 (1.405) | 0.458 (0.118) | 2.332 (1.951) | 77.613 (8.242) |
|  |  | 0.033 - 0.997 | 2-8 | 0.086 - 0.689 | 0.782 - 16.730 | 62.500 - 106.000 |
|  | *Ctenomys lami* | 0.377 (0.377) | 3.172 (2.272) | 0.328 (0.293) | 1.597 (1.191) | 76.291 (9.325) |
|  |  | 0.043 - 1.858 | 2-15 | 0.293 - 1.809 | 0.382 - 8.451 | 57.800 - 107.700 |
| Squeal | *Ctenomys torquatus* | 0.125 (0.125) | 1.400 (0.632) | 0.477 (0.172) | 1.276 (0.564) | 76.893 (8.212) |
|  |  | 0.044 - 0.533 | 1 - 3 | 0.172 - 0.775 | 0.778 - 2.659 | 64.200 - 92.000 |
|  | *Ctenomys lami* | 0.104 (0.169) | 1.275 (0.507) | 0.416 (0.246) | 1.441 (0.763) | 71.317 (9.422) |
|  |  | 0.022 - 1.344 | 1 - 3 | 0.086 - 1.206 | 0.496 - 4.721 | 51.800 - 100.800 |
| Squeal - Subtype |  | 0.354 (0.376) | 2.200 (1.867) | 0.666 (0.224) | 2.212 (0.800) | 71.947 (9.520) |
|  |  | 0.070 - 1.257 | 1 - 7 | 0.086 - 0.948 | 1.343 - 3.689 | 58.100 - 93.800 |
| Sequential | *Ctenomys torquatus* | 1.904 (1.377) | 23.883 (25.104) | 0.172 (0.155) | 0.748 (0.389) | 85.152 (8.401) |
|  |  | 0.198 - 8.281 | 2 - 147 | 0.086 - 0.603 | 0.333 - 2.154 | 70.400 - 102.100 |
|  | *Ctenomys lami* | 2.266 (2.452) | 15.944 (14.534) | 0.233 (0.178) | 1.424 (1.075) | 88.768 (14.580) |
|  |  | 0.250 - 19.000 | 2 - 83 | 0.086 - 0.861 | 0.390 - 7.845 | 64.200 - 133.800 |

## Description of the identified signal types

### Mono

It is a pulsed signal formed by a single low frequency note (Fig 3).

**Fig 3.**
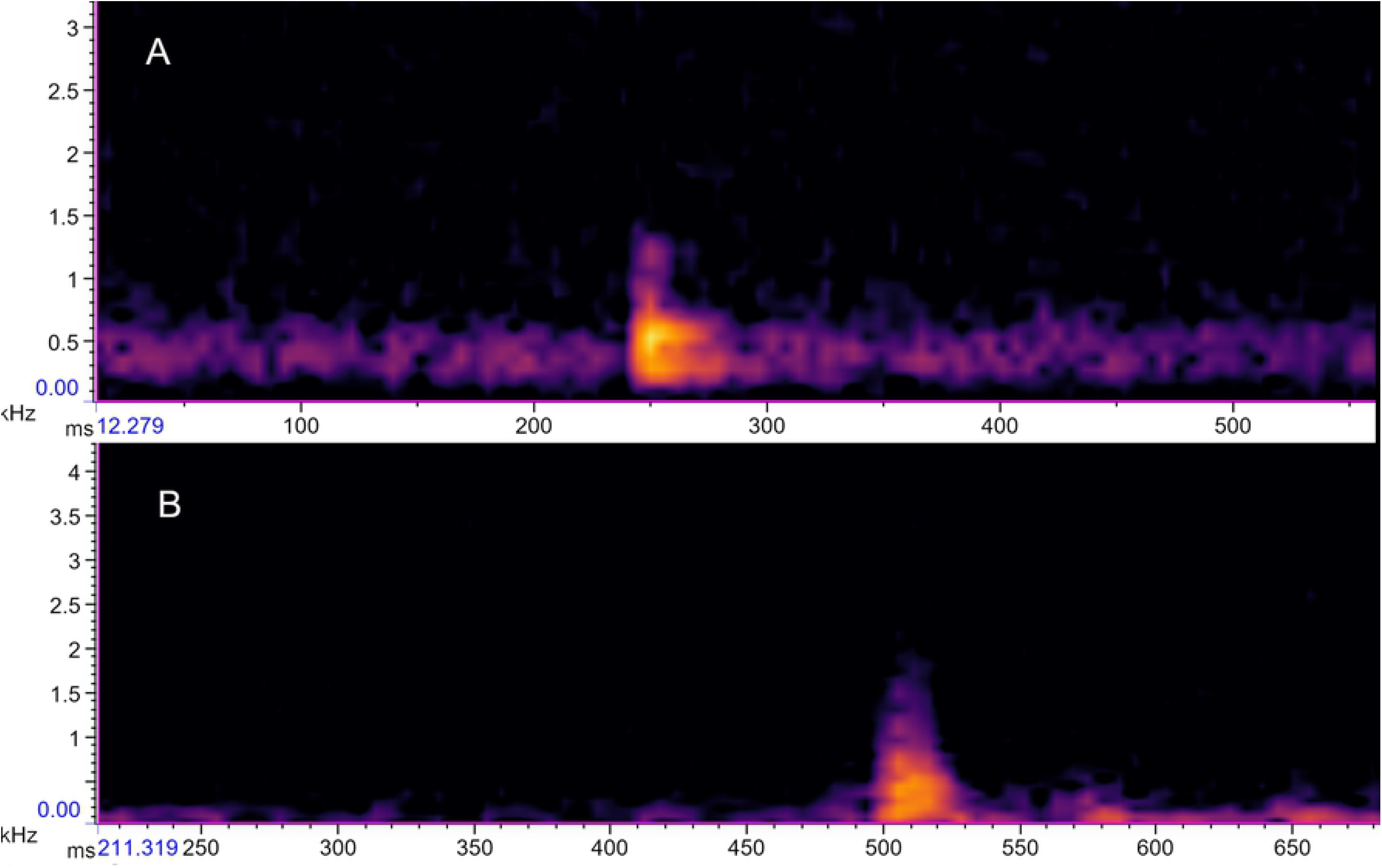
Mono signal. Single note spectrogram of the mono signal in *C. torquatus* (A) and *C. lami* (B).

For *C. torquatus* it was possible to identify 122 signals (Fig 2) present in 7 individuals, emitted by females, males and unidentified individuals (Table 1). In *C. lami*, 90 signals (Fig 2) were found in 10 individuals, emitted by females and unidentified individuals (Table 2). The mean values of the acoustic parameters investigated in the mono signal of both species are shown in Table 3.

### Drummed

It is a pulsed signal, analogous to a non-vocal sound. Formed by serial notes, with a silence interval between the notes, constituting the drummed phrases (Fig 4).

**Fig 4.**
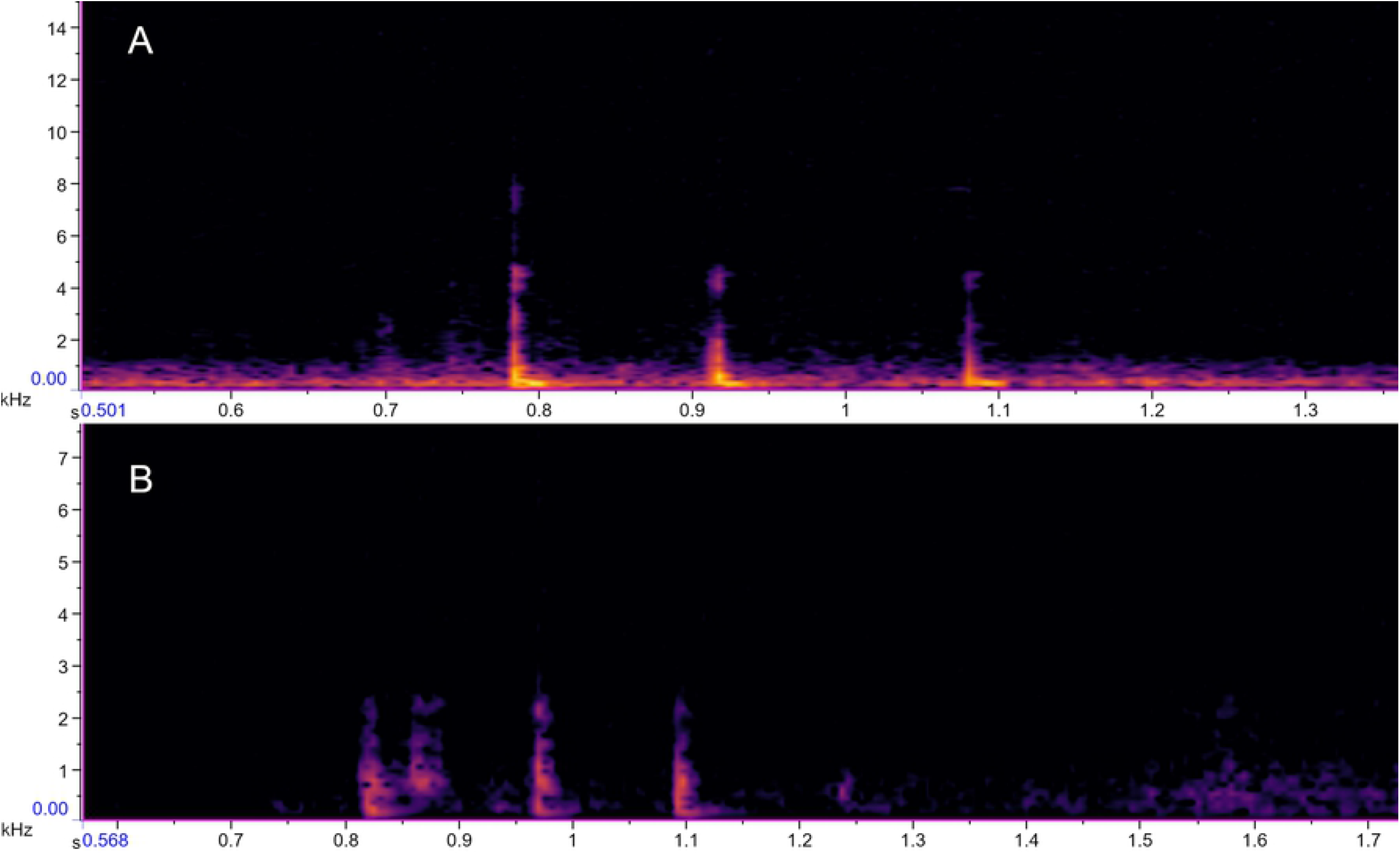
Drummed type signal. Spectrogram of the drummed phrase with 3 notes in *C. torquatus* (A) and 3 notes in *C. lami* (B).

For *C. torquatus,* 115 drummed signals (Fig 2) emitted by 9 individuals were identified. The phrases ranged from 2 to 8 notes, emitted by females, males, and unidentified (Table 1). In *C. lami,* the drummed signal comprised phrases from 2 to 15 notes, being distinguished 64 signals (Fig 2) of this type registered for 14 individuals, emitted by females and unidentified individuals (Table 2). The mean values of the acoustic parameters investigated in the drummed signal of both species are shown in Table 3.

### Tuc

They are low frequency signals, formed by a set of notes with a short interval of silence separating one note from another, thus forming the tuc phrases (Fig 5). The sign tuc is an onomatopoeia that originated the popular name of the animal: tuco-tuco.

**Fig 5.**
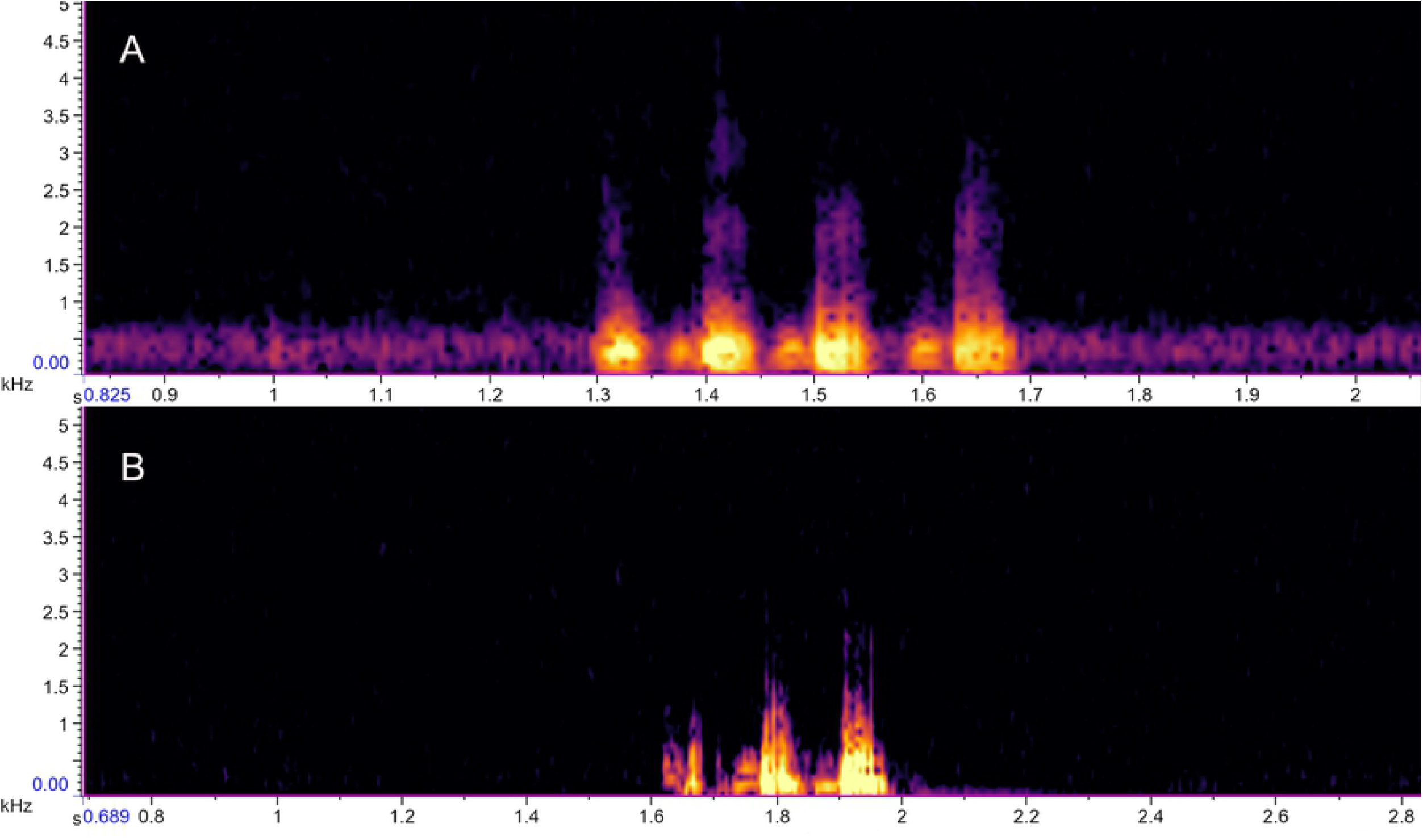
Tuc signal. Spectrogram of the tuc phrase with 4 notes in *C. torquatus* (A) and 3 notes in *C. lami* (B).

In *C. torquatus,* it was the signal that showed the most incidence, being recognized 474 tucs phrases (Fig 2). In only 1 of 14 recorded individuals, it was not possible to identify this signal. The phrases’ length ranged from 2 to 12 notes, emitted by females, males, and unidentified individuals (Table 1). For *C. lami,* 119 tuc phrases were identified (Fig 2). The number of notes in the phrases ranged from 2 to 12 notes. The tuc type signal was not verified in only 1 of the 15 selected individuals, being emitted by females and unidentified individuals (Table 2). The mean values of the acoustic parameters investigated in the tuc signal of both species are shown in Table 3.

### Sequential

It is a rhythmic signal, with similar notes in continuity separated by a short silence, constituting the phrases (Fig 6).

**Fig 6.**
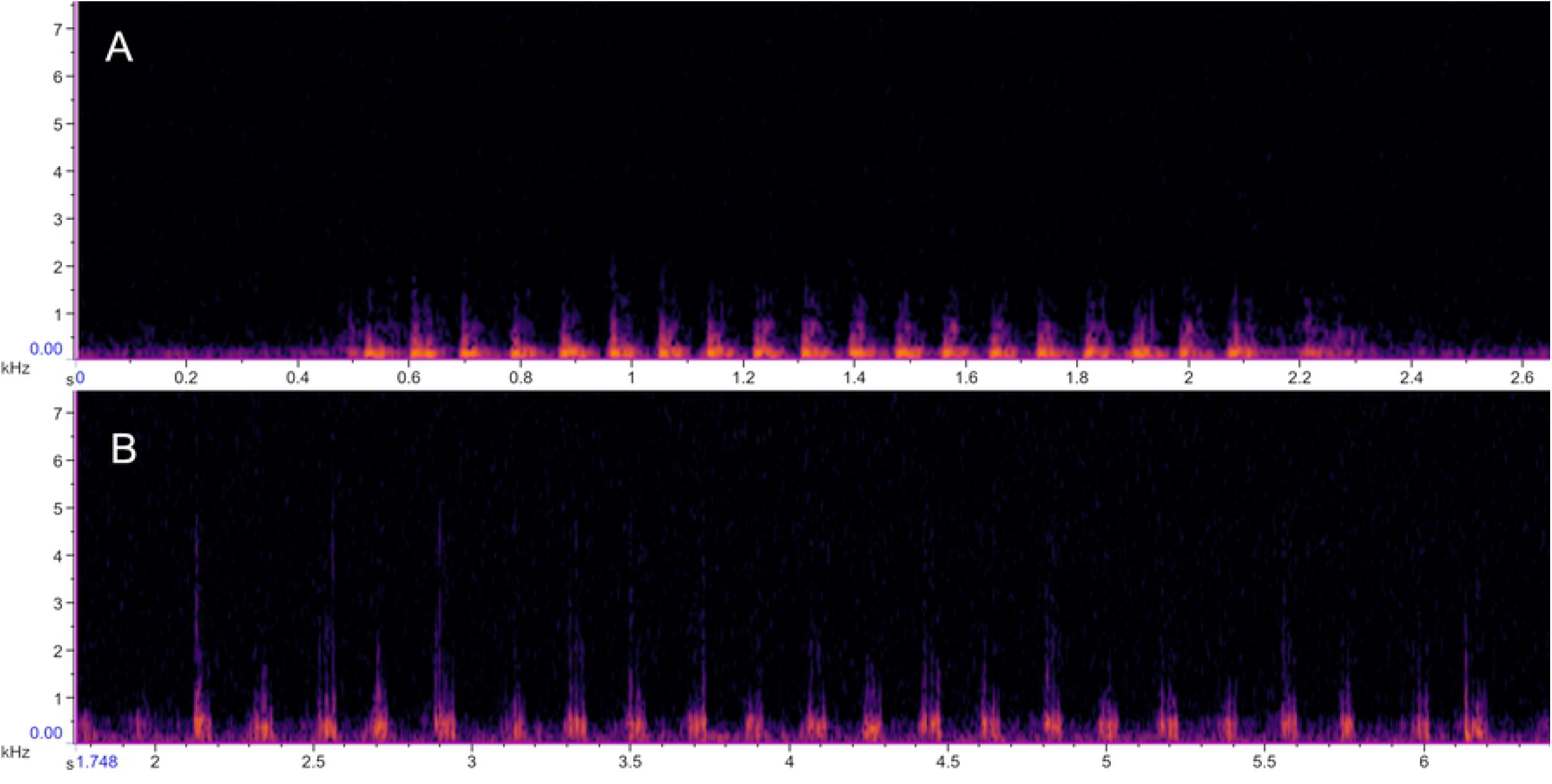
Sequential type signal. Spectrogram of the sequential phrase with 20 notes in *C. torquatus* (A) and 22 notes in *C. lami* (B).

For *C. torquatus*, 60 sequenced phrases (Fig 2) were recognized, emitted by 6 individuals, females, males, and unidentified (Table 1). In *C. lami,* it was the signal that showed the most incidence, totaling 197 phrases (Fig 2) sequenced and expressed by 13 individuals, females, and unidentified (Table 2). The mean values of the acoustic parameters investigated in the sequential signal of both species are shown in Table 3.

### Squeal

It is a harmonic signal and can be emitted in single note or in serial form. In serial form the notes are separated by a silence interval, generating the squeal phrases (Fig 7).

**Fig 7.**
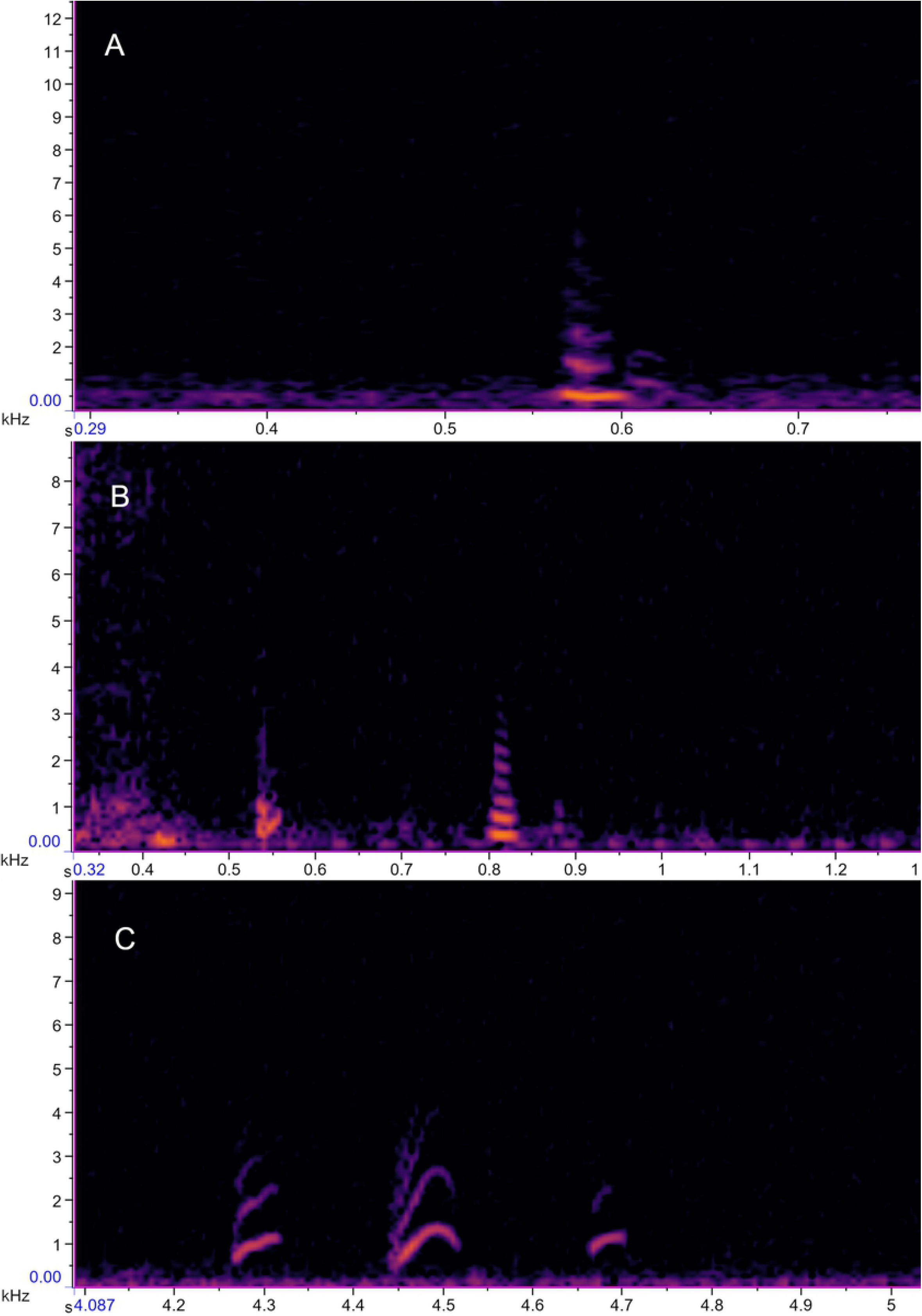
Squeal type signal. Squeal phrase spectrogram with 1 note in *C. torquatus (A)* and 2 notes in *C. lami* (B). And the 3 notes squeal subtype (C) identified in *C. lami*.

In *C. torquatus,* the squeal phrases ranged from 1 to 3 notes, totaling 15 phrases (Fig 2) registered in 7 individuals: females, males, and unidentified (Table 1). For *C. lami*, 109 squeal phrases were identified (Fig 2), ranging from 1 to 3 notes, emitted by 12 female and unidentified individuals (Table 2). The mean values of the acoustic parameters investigated in the squeal signal of both species are shown in Table 3. A squeal subtype was identified in *C. lami* (Fig 7 C), a signal with a more stressed harmonic. 15 phrases were recognized, emitted by 3 individuals: female, and unidentified (Table 2), the phrases presented from 1 to 7 notes. The mean values of the acoustic parameters investigated in the squeal subtype are shown in Table 3.

## Discussion

It was possible to verify, in both species, a sharing of the investigated sound signals, expressing similarity in both acoustic parameters analyzed and in the spectrographic design (Table 3) (Fig 3, 4, 5, 6, and 7). These analogies may be related to the similarity of environments that these species occupy [34,35] or to the fact that they belong to the same phylogenetic group, Torquatus [37].

According to the mean data obtained from the frequency bands used (Table 3), most of the sequences presented low frequency, which is expected for an subterranean animal, due to limiting factors found in the structuring of the environment the animal is inserted [23,24]. The improving of the middle ear cavities of these animals is also associated with hearing low-frequency signals [46]. In addition, low-frequency signals have been reported for other *Ctenomys* species such as *C. pearsoni, C. talarum,* and Anillaco *Ctenomys* sp. [19,20,25].

One of the reasons why *C. torquatus* and *C. lami* commonly use low-frequency signals is that high-frequency sounds are very directional in open environments, being relevant in contact or agonistic calls, however, when emitted in an underground environment, they can be promptly dispersed within the tunnels thus impairing communication, especially over long distances [30].

Some studies have already been performed to verify the propagation of sounds within the burrows. The propagation of calls of mole rats *(Spalax ehrenbergi)* at different frequencies was tested. and it was observed that 440 Hz signals propagated more efficiently over short distances within the burrows [47]. Lange et al. [23] evaluated the transmission of sounds with different types of frequencies in *Fukomys mechowii* and *Fukomys kafuensis.* Although the burrows had different diameters, the sound propagation was very similar between them. Low-frequency sounds undergo less attenuated than high-frequency sounds and still had their amplitudes increased, called the “stethoscope” effect [23]. In *C. talarum* [24], the relationship between the effectiveness of signal propagation and the entrance structure of the burrows was investigated, but in a similar system to natural burrows. The “stethoscope” effect was observed as in *F. mechowii* e *F. kafuensis* [23].

Sound propagation can be affected by many factors [1], however, in different subterranean species, the acoustic attributes present inside the tunnels are similar, which could have led to a convergent adaptation in the bioacoustics of subterranean rodents [48]. Studies that investigate acoustic communication can act as an auxiliary tool in the interpretation and explanation of general evolutionary concepts.

The mono signal was frequently detected in individuals of both species, as a pulsed signal, with a single note and low frequency (Table 3). Single note signals could play a part in a longer vocal phrase. The emitting of only part of the calling could be related to the motivational state of the emitter individual [2,4].

The rhythmic patterns of the calling emitted can be related to the information in the signals [17,26]. As a pulsed single note signal generally does not variate in the rhythmic pattern or possibilities of alternating the emission’s arrangement, it is a single note phrase.

Eventually, this type of signal could not convey information to another individual receiver, but a reference to the emitter itself.

The mono signal may act as an echolocation instrument, along with other sensory channels present in these animals [49]. Mechanisms of cognition have been proposed from the correlation with spatial orientation and spatial navigation to rodent sensory systems [50]. It has been observed that *Spalax ehrenbergi* uses seismic signal reflection to detect objects, which can be considered a way of echolocating [51].

Tuco-tucos may use the stethoscope effect of tunnels [23] to amplify the propagation of the sent pulsed signal and use the echoed signal, which would conserve enough energy to be decoded by the transmitter, as a way of locating within the tunnels. There are no reports of echolocation in *Ctenomys.* As this is a field that had never been addressed, more precise investigations are necessary before confirming the hypothesis.

The drummed signal was the one that sounded different. A uniform signal with a noise similar to the touch of two objects. Mechanical sounds, such as teeth chattering, have already been registered in *Ctenomys talarum* [20]. The chattering of teeth was described as an emission of short and quick notes, recorded during the feeding process and agonistic contexts among males [20].

Studies are describing that subterranean rodents can produce signals beyond vocalizations, called seismic signals [21,22,52–57]. These signals can be generated from the behavior of stomping feet [53,57–60] and/or head [22,52,54,61] inside the tunnels as a way to produce noise and use it to communicate. The drummed signal may be the result of this kind of sound production. However, we cannot confirm this assumption since we did not have visual access to the behavior performed by the animals.

Depending on the frequency range and the volume used in a call, the sound signals can travel long distances and can be considered one of the main forms of communication between species that live in the underground environment [23,24,46]. In *Ctenomys,* long range signals have been described as high-volume, low-pitched, and repetitive note signals [19,20,25]. Two patterns for long-range signals have been recognized until this date [33]. In *C. torquatus* and *C. lami,* the tuc and sequential signals showed characteristics of long-range signals [19,20,25] (Table 3).

The tuc type vocalizations are mostly emitted in isolation and with variable repetition in the number of notes (Table 3). The tuc signal has already been characterized for *C. pearsoni, C. talarum* and Anillaco *Ctenomys* sp, named in these species as: S-type – *C. pearsoni* [19], tuc-tuc – *C. talarum* [20] and as LRV (long-range vocalizations) in Anillaco *Ctenomys* sp [25]. Sonically, according to the authors’ description, the tuc type signal emitted by *C. torquatus* and *C. lami* is similar to the description of these referred signals, but they have distinct morphological and acoustic arrangement characteristics in the spectrographic design. However, the low sound frequency range is shared between species (Table 4).

**Table 4.** Means of dominant frequencies (MDF) of tuc signals and their equivalents.

| Species | Signal | MDF |
| --- | --- | --- |
| *C. torquatus* | Tuc | 0,220 |
| *C. lami* | Tuc | 0,219 |
| *C. pearsoni* | S | 0,243 |
| *C. talarum* | Tuc-tuc | 0,257 |

MDF = Means of dominant frequencies in kHz.

The tuc-equivalent signals in *C. talarum, C. pearsoni,* and Anillaco *Ctenomys* sp. are recognized as long-distance signals, which reiterates the character of the tuc as a long-range signal.

The sequential type signal may also be characterized as long-distance because it is analogous to the physical description made for long-range signals [33]. The same way it resembles the mating signal emitted by male individuals of *C. talarum* [20], however in *C. torquatus,* it was emitted by both sexes (Table 1), and in *C. lami* it was possible to confirm only the emission by females since all animals captured and identified were females (Table 2). In addition to morphological similarity, they also use an approximate frequency range during emissions (Table 5).

**Table 5.** Means of the dominant frequencies of sequential signals and their equivalents.

| Species | Signal | MDF |
| --- | --- | --- |
| *C. torquatus* | Sequential | 0,172 |
| *C. lami* | Sequential | 0,233 |
| *C. talarum* | Tuc-tuc | 0,257 |
| *C. talarum* | Mating | 0,400 |

MDF = Means of dominant frequencies in kHz.

The attributes observed in the tuc and sequential signals suggest that they may present characteristics of territorial and mating signals, as both would need to travel long distances to find the required potential receptors. The variability in the ordering of emitted signals may be related to the motivational state of the emitting individual [2,4], and the information encoded in the sound signals may be associated with the rhythmic pattern of the observed emissions [17,26].

In contrast, the squeal signal presented a relatively high-frequency range compared to the values of the other signal types described (Table 3). In many mammals, the emission of high-pitched signals may be related to the calls of pups [4]. Acoustic signals produced by puppies have already been described for *C. talarum* [29] and *C. pearsoni* [30]. These puppies’s calls show a generally higher frequency range than the other calls and exhibited characteristics of modulated signals [29,30]. The squeal calls of *C. lami* and *C. torquatus* also have these attributes (Fig 4). Morphologically, the spectra of squeal phrases are similar to the structure of some signals described for *C. pearsoni* and *C. talarum* infants [29,30]. In *C. lami,* the squeal subtype presents a very prominent modulation and analogous to the care calls of *C. talarum* and *C. pearsoni* [29,30].

*C. lami* emitted a considerably larger amount, and greater sound diversity of squeal call than *C. torquatus,* which may be justified because of recordings were made with *C. lami* just after the reproductive period of this species [35], reinforcing the assumption that puppies or even juveniles may have emitted squeal phrases.. Though a more detailed investigation of this call is necessary, especially in *C. torquatus* in the post-reproductive season.

In general, the distinction between the characteristics of the signals of *C. torquatus* and *C. lami* with the other species already studied, both in spectrographic morphology and in the physical attributes of the acoustic signals investigated in this work, maybe the result of a geographical variation [31,62], speciation events [63–65], ecological factors [66,67], physical characteristics of the environment [1] or the motivational state of the emitters [4]. However, it is necessary to conduct more specific studies focusing on answering these questions. The essence of this work was to describe the acoustic signals in which emission was captured in the field and to understand the physical-morphological attributes of these signals.

By this, the characterization of the present type of sound signals used by *C. torquatus* and *C. lami* in the field promotes access to the main acoustic parameters used by these animals during the emission of signals. This open new perspectives for in-depth comparative analyzes and studies investigating the influence of anthropogenic noises on the vocal communication of these species.

This study was carried out based on the development of an incipient methodology, as bibliographically registered work on a methodology for sound capture of subterranean rodents inside their burrows in a natural environment was not found. Until recent times there was difficult making direct recordings into the tunnels, notably the underground environment presents its difficulties. According to the methodology applied, we have a boundary of reliability that the signals emitted inside the tunnels were fully captured. The methodology used is open to improvement, however it was able to accomplish one of the main purposes of obtaining organic vocalizations of the tuco-tuco interfering as little as possible in the animal and its habitat.

## Author Contributions

Conceptualization: Keila C. Zaché, Lucas Machado Silveira, Gabriel Francescoli, Thales

Renato Ochotorena de Freitas

Data Curation: Keila C. Zaché

Formal Analysis: Keila C. Zaché

Investigation: Keila C. Zaché, Lucas Machado Silveira

Methodology: Keila C. Zaché

Writing – Original Draft Preparation: Keila C. Zaché

Writing – Review & Editing: Keila C. Zaché, Lucas Machado Silveira, Gabriel Francescoli, Thales Renato Ochotorena de Freitas

## Acknowledgments

The authors would like to thank the team at the Estação Ecológica do Taim and the Reserva Biológica do Lami José Lutzenberger for authorizing the sampling, for supporting our work and making the research feasible. We also thank Thamara de Almeida and Ana Maria O. Mastella for sampling assistance in some fieldworks.

